# Production of spliced long noncoding RNAs specifies regions with increased enhancer activity

**DOI:** 10.1101/319400

**Authors:** Noa Gil, Igor Ulitsky

## Abstract

Active enhancers in mammals produce enhancer RNAs (eRNAs), that are bidirectionally transcribed, unspliced, and unstable noncoding RNAs. Enhancer regions are also enriched with long noncoding RNA (lncRNA) genes, which are typically spliced and are longer and substantially more stable than eRNAs. In order to explore the relationship between these two classes of RNAs and the implications of lncRNA transcription on enhancer functionality, we analyzed DNAse hypersensitive sites with evidence of bidirectional transcription, which we termed eRNA producing centers (EPCs). A subset of EPCs, which are found very close to the transcription start site of lncRNA genes, exhibit attributes of both enhancers and promoters, including distinctive DNA motifs and a characteristic landscape of bound proteins. These EPCs are associated with a subset of relatively highly active enhancers. This stronger enhancer activity is driven, at least in part, by the presence of evolutionary conserved, directional splicing signals that promote lncRNA production, pointing at a causal role of lncRNA processing in enhancer activity. Together, our results suggest a model whereby the ability of some enhancers to produce lncRNAs, which is conserved in evolution, enhances their activity in a manner likely mediated through maturation of the associated lncRNA.

## Introduction

Enhancers are DNA regulatory elements that can activate transcription of distally located genes, typically acting in a spatially and temporally restricted manner. Active enhancer regions are demarcated by distinct chromatin marks such as H3K27ac and H3K4me1, bind CBP/p300, and overlap DNAse hypersensitive sites (DHSs) (Calo and Wysocka 2013). Enhancers are thought to serve as platforms for the assembly of transcription factors (TFs) and the Pol II preinitiation complex, signals which are then relayed to the promoters of target genes through chromatin loops that can be either pre-formed or induced upon enhancer activation, resulting in increased activity of the promoter and increased gene expression (Shlyueva, Stampfel, and Stark 2014; Blackwood and Kadonaga 1998).

This model of enhancer activity dictates that active enhancers are commonly bound by factors required for initiating transcription. High-throughput sequencing has accordingly unveiled extensive transcription emanating from enhancer elements, the products of which are referred to as enhancer RNAs (eRNAs) (De Santa et al. 2010; Kim et al. 2010; Koch et al. 2011; Hah et al. 2013). Pol II activity and bidirectional eRNA production are now considered a hallmark of active enhancers, and are being increasingly used to annotate such elements in the genome (Melgar, Collins, and Sethupathy 2011; Nagari et al. 2017). eRNAs are typically relatively short (~1-3Kb on average) and unspliced (De Santa et al. 2010; Kim et al. 2010). They are also unstable, and do not accumulate to substantial levels in cells (Djebali et al. 2012; De Santa et al. 2010). As eRNAs are not readily detectable in steady-state RNA-seq data, their annotation relies on sequencing of nascent RNA or on perturbations of nuclear decay pathways (Hah et al. 2011; Robin Andersson, Refsing Andersen, et al. 2014; Pefanis et al. 2015; Core et al. 2014; Austenaa et al. 2015). However, this definition is not all-inclusive, as some eRNAs are produced unidirectionally and are polyadenylated (Koch et al. 2011), and are therefore presumably more stable. Several roles have been proposed for eRNAs. These include promoting the formation of chromatin loops between the enhancers and their target promoters (Hsieh et al. 2014; Li, Lam, and Notani 2014), or acting to increase transcription at promoters after such loops have been formed, for example by remodeling chromatin or promoting Pol II elongation (Mousavi et al. 2013; Schaukowitch et al. 2014). It has also been suggested that the act of eRNA transcription at enhancers, rather than the mature RNA product, might be important for enhancer function (Natoli and Andrau 2012).

An additional species of non-coding RNAs (ncRNAs) that has recently received much attention are long non-coding RNAs (lncRNAs). Unlike eRNAs, lncRNAs are polyadenylated and typically spliced (Ulitsky and Bartel 2013), and therefore constitute more stable transcripts. Several genome-wide annotation studies identified an enrichment of lncRNA genes in the vicinity of enhancers, with up to 30–60% of lncRNAs overlapping regions with enhancer characteristics (Vučićević et al. 2015; Werner and Ruthenburg 2015; De Santa et al. 2010). In parallel, focused studies of specific lncRNAs found that they act to increase expression of genes in *cis*, functions which are possibly related to enhancers found in the vicinity of the lncRNAs, though the nature of the relationship and the mechanism by which these lncRNAs function remain mostly unclear (Isoda et al. 2017; Anderson et al. 2016; Engreitz et al. 2016; Lai et al. 2013). Importantly, some lncRNAs produced from enhancer-containing loci have also been proposed to act in a similar manner to eRNAs (Ørom et al. 2010; Lai et al. 2013; Xiang et al. 2014), emphasizing that these two categories of transcripts are not mutually exclusive. Accordingly, it is likely that some of the unidirectional, stable eRNA transcripts are de facto lncRNAs (Natoli and Andrau 2012), and it is unclear whether the differences between the two groups of ncRNAs confer any distinct functions. The promoters of some lncRNAs and PCGs were shown to have enhancer activity (Engreitz et al. 2016; Dao et al. 2017; Nguyen et al. 2016), further confounding the relationship between enhancers and promoters.

Taken together, it appears that the transcriptional landscape at enhancers is complex and produces an array of non-coding transcripts which differ in their structural characteristics and stability; it is presently unclear what, if any, are the functional distinctions between these classes of ncRNAs, nor what are the consequences of production of stable transcripts on enhancer activity. In this study, we set out to compare enhancers that produce different types of ncRNAs. We begin by showing that while some enhancers produce only eRNAs, others produce both eRNAs and lncRNAs. The ability of some enhancers to drive the production of lncRNAs is correlated with a generally higher enhancer activity, which is apparently mediated by the recruitment of RNA binding proteins and specifically splicing factors, presumably required for the formation of the mature lncRNA transcript.

## Results

### Chromatin accessibility and bidirectional transcription define enhancer regions

In order to study the relationship between eRNAs and enhancer-associated lncRNAs, we first set out to annotate eRNAs from publicly-available GRO-seq datasets in four cell lines: the human ENCODE cell lines K562, HepG2, and MCF7, and mouse embryonic stem (mES) cells (see Methods for data sources). Enhancers can be annotated using various hallmarks, including chromatin marks, DHSs, CAGE data, and various RNA sequencing modalities; after experimenting with various approaches and published pipelines, we found that DHSs offer the best resolution for eRNA annotations along with relatively high sensitivity. We therefore searched for evidence of substantial bidirectional transcription stemming from regions surrounding DHSs in each of the four cell lines. For each pair of bidirectional transcripts, we identified the center coordinate (the coordinate which offered the best separation between the GRO-seq signal on the two strands; see Methods, Figure S1A), retaining only bidirectional transcripts remote from PCGs (Figure 1A). We termed these positions eRNA-producing centers (EPCs). The EPCs detected by our scheme overlap the majority of, and substantially outnumber, the eRNAs detected by the groHMM algorithm (Chae, Danko, and Kraus 2015) (Figure S1B), show canonical enhancer characteristics such as high H3K4me1 and H3K27ac signal, and tend to overlap active chromatin regions annotated by ENCODE combined segmentations (Hoffman et al. 2013) (Figures 1B, S1C,D). We annotated 4,470–15,244 EPCs in each of the four cell lines (**Table S1**). When examining regions ±1Kb from the EPCs, 64% and 67% of K562 and HepG2 EPCs, respectively, overlap a chromatin region annotated as an enhancer in these cell lines by ENCODE combined segmentations (Hoffman et al. 2013).

**Figure 1.**
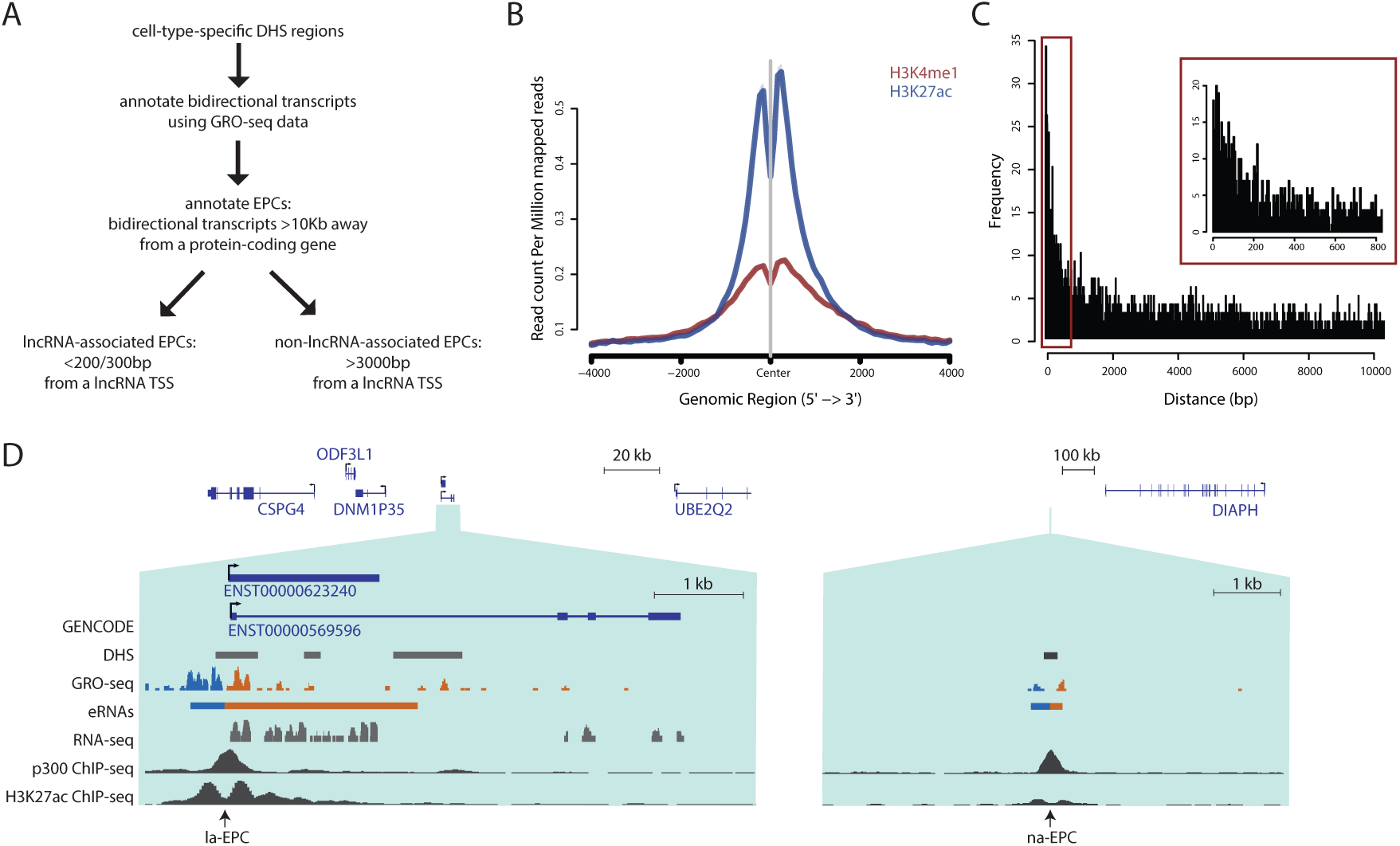
EPC annotations. **A**. eRNAs were annotated by detecting bidirectional GRO-seq signal from DHS regions that are >10Kb away from protein-coding genes. The ‘center’ of the eRNA pair, or the EPC, was defined as the position at which transcription to either side provides the best separation between reads mapping to opposite strands. EPCs were divided into lncRNA-associated EPCs (la-EPCs) or non-lncRNA-associated EPCs (na-EPCs) based on the distance of the EPC to the closest GENCODE lncRNA TSS. **B**. Metagene plots showing the distribution of H3K27ac and H3K4me1 ChIP-seq coverage signals around K562 EPCs. **C**. Distribution of distances between K562 EPCs and the closest lncRNA TSS. Region in the red rectangle is shown with higher bin resolution in inset. D. Genome browser images with examples of an la-EPC (left; hg19 coordinates of enlarged region chr15:76050001-76056500) and an na-EPC (right; hg19 coordinates of enlarged region chr13:60039501-60046000). Tracks in both images are normalized to the same scale. Only representative GENCODE transcript models are shown. GRO-seq data is the same as used for EPC annotations, and color indicates the strand to which the reads mapped. DHS, RNA-seq, and p300 and H3K27ac ChIP-seqs were all taken from ENCODE K562 data.

### A subset of EPCs serve as TSSs for lncRNA production

We next examined the distribution of distances between those fine-mapped EPCs and the nearest GENCODE lncRNA TSS (Figure 1A), and found a group of EPCs that are in close proximity to lncRNA TSSs (Figures 1C and S1E). Accordingly, we classified EPCs into lncRNA-associated EPCs (la-EPCs; 3–5% of all EPCs) and non-lncRNA-associated EPCs (na-EPCs; 81–88% of all EPCs), based on the distance between the EPC and the nearest lncRNA TSS (see Methods; **Table S1**; Figures 1A,C,D and S1E). la-EPCs, but not na-EPCs, are associated with strong poly(A)+ RNA-seq signal (Figures 2A and S2A), and 43–67% of annotated lncRNAs that are associated to an EPC are expressed (FPKM>0.2) in these cell lines. This suggests that the bidirectional transcription that originates from some enhancers is elongated unidirectionally to form a stable lncRNA transcript. The transcription to the other side of the stable lncRNA transcript is reminiscent of the similarly unstable and unspliced Promoter Upstream Transcripts (PROMPTs) transcribed divergently to PCGs (Preker et al. 2011; Flynn et al. 2011; Ntini et al. 2013), highlighting the role that transcribed enhancers can play in the evolution of new lncRNA genes by contributing novel TSSs (see Discussion).

**Figure 2.**
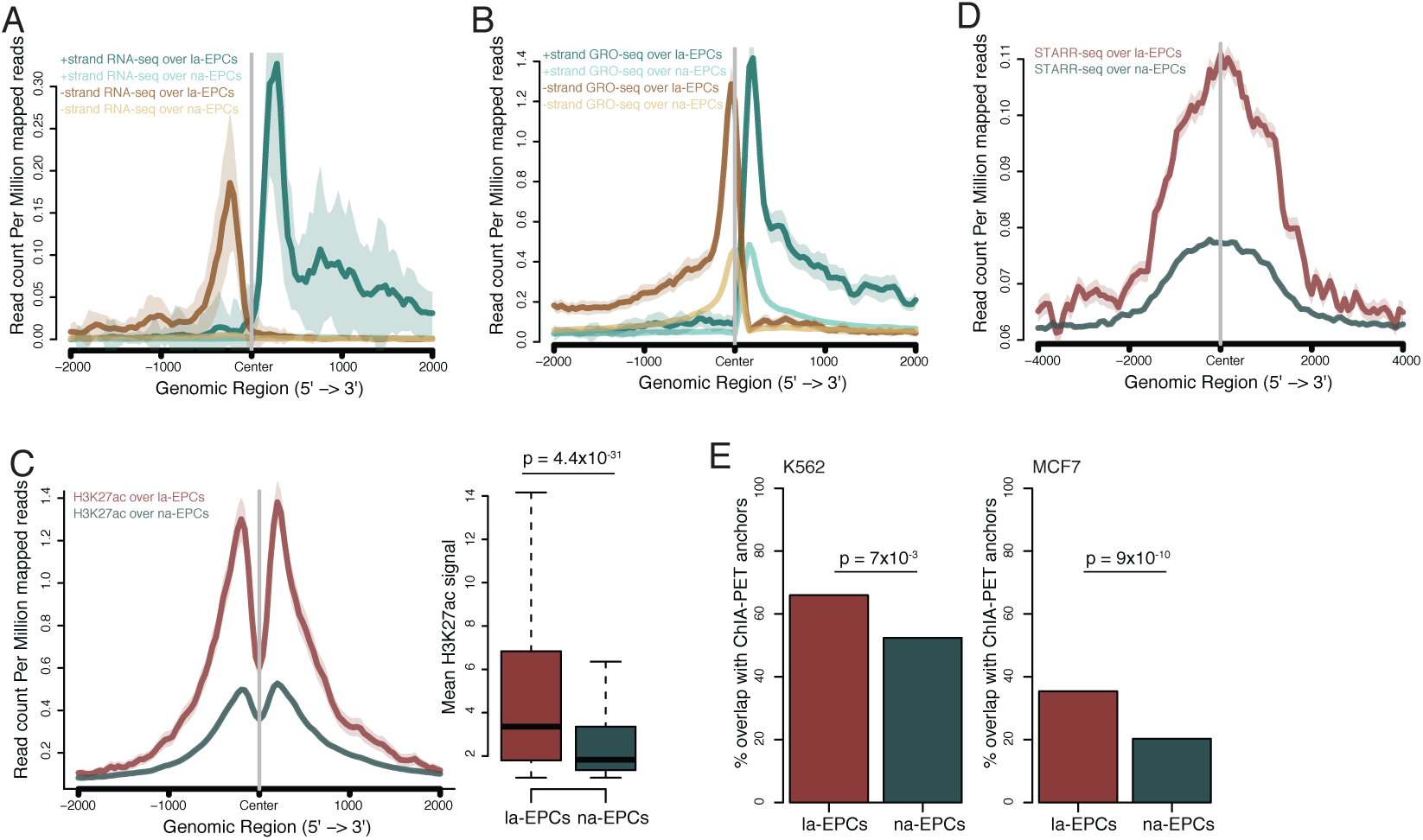
Differences between la-EPC and na-EPC loci. **A**. Metagene plots of K562 RNA-seq read coverage, separated by strand, over la-EPCs and na-EPCs. Shaded regions represent standard errors. **B**. As in A, for GRO-seq signal. **C**. Left, metagene plot of K562 H3K27ac ChIP-seq signal over la-EPCs and na-EPCs; right, quantification of H3K27ac signal over the region ±1Kb around the EPCs. P value calculated using two-sided Wilcoxon test. Shaded regions in the left plot represent standard errors. **D**. Metagene plot of HeLa-S3 STARR-seq over la-EPCs and na-EPCs combined from the three human cell lines. Shaded regions represent standard errors. **E**. Percent of la-EPCs or na-EPCs which overlap Pol II ChIA-PET anchors in the indicated cell lines. P values calculated using the proportion test.

While both la- and na-EPCs are associated with peaks of coverage of GRO-seq and H3K27ac ChIP-seq reads, both these marks, associated with enhancer strength, are significantly stronger in la-EPCs (Figures 2B,C and S2B,C), suggesting that enhancers associated with lncRNA production are more active than other enhancers. This increased enhancer activity is intrinsically encoded in the sequences of la-EPC regions, as la-EPCs also exhibit a higher STARR-seq signal, which measures enhancer activity in a non-native context (Muerdter et al. 2018) (Figure 2D). Further, la-EPC loci overlap significantly more Pol II ChIA-PET anchors than na-EPCs (Figures 2E and S2D), indicating that the enhancers they overlap are more likely to be bound by Pol II and form spatial contacts with distal loci; similarly, mES la-EPCs also overlap more YY1 ChIA-PET peaks (and CTCF, to a lesser extent) (Figure S2D), a protein involved in the maintenance of enhancer-promoter loops (Weintraub et al. 2017). Together, these findings suggest that la-EPCs comprise a subgroup of enhancers associated with stronger enhancer activity than na-EPCs.

### lncRNA-associated EPCs have a distinct chromatin and protein binding landscape

The apparent differences between la-EPCs and na-EPCs may result from differences in the chromatin modifications or the proteins binding the enhancer region and driving Pol II recruitment and/or transcriptional elongation. Therefore, we next assayed for differential protein occupancy at la-EPCs vs. na-EPCs by examining ChIP-seq peaks from available ENCODE datasets and comparing the number of binding peaks in la-EPCs and in random samplings of na-EPCs matched to la-EPCs in their H3K27ac coverage. As expected for regions associated with more substantial transcription initiation, a higher percentage of la-EPCs in all four cell lines are associated with Pol II peaks, as well as the promoter mark H3K4me3 (Figure S3A), further demonstrating that these enhancers also serve as promoters of lncRNA genes. This is in agreement with previous studies showing that lncRNA promoters show both enhancer and promoter characteristics (Marques et al. 2013; Lam et al. 2014; R. Andersson, Sandelin, and Danko 2015). Importantly, CTCF is one of the proteins enriched in la-EPCs in all human cell lines (but not in mES; data not shown), further demonstrating that la-EPC loci are more involved in chromatin looping than na-EPCs (Figures 3A and 2D).

**Figure 3.**
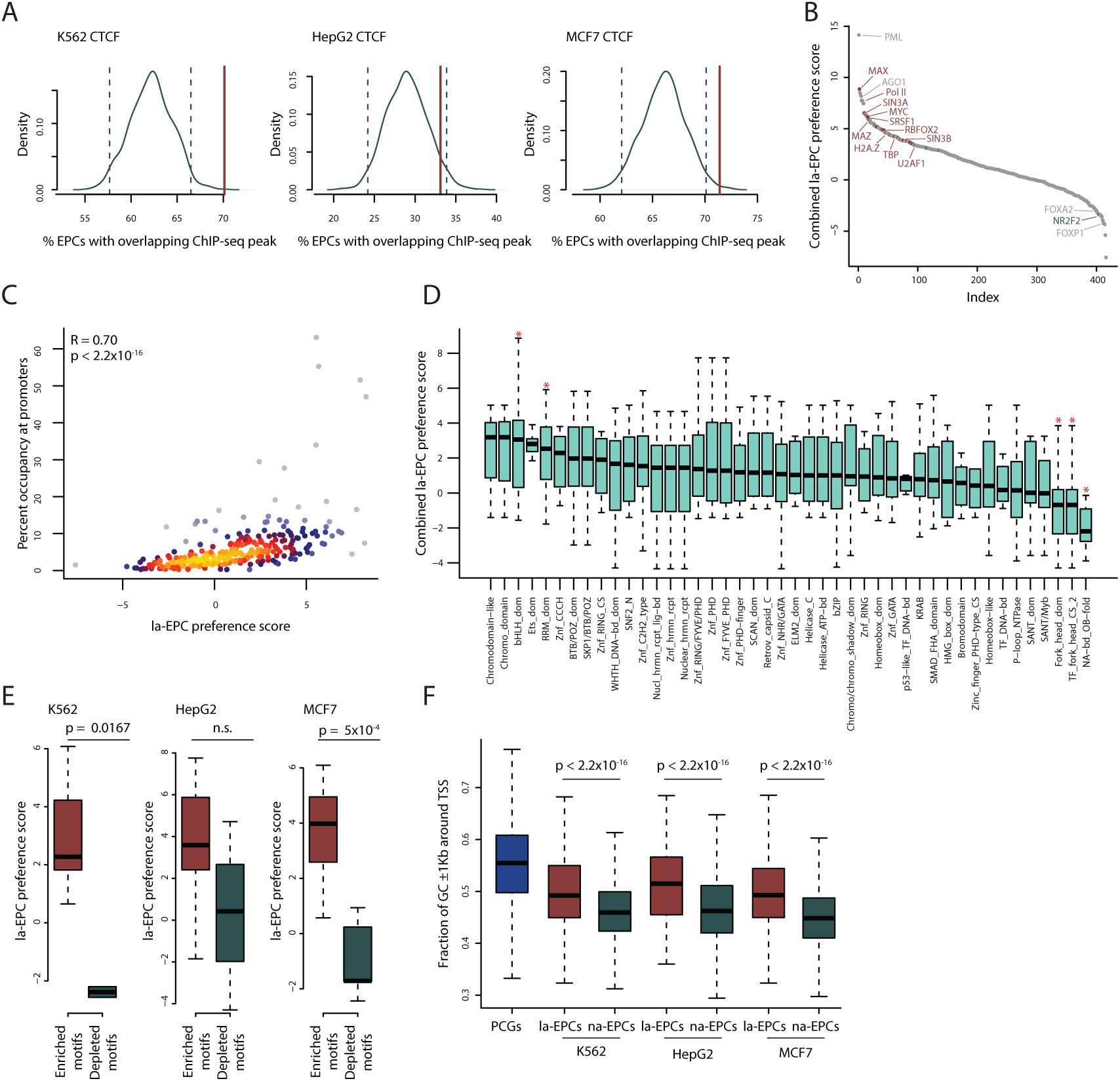
Differences in protein binding patterns at la-EPC and na-EPC loci. **A.** Green lines: Distribution of the percent of na-EPC loci with overlapping CTCF ChIP-seq peaks in a random sampling of na-EPC loci in the indicated cell lines. Dashed lines: 2.5^th^ percentiles of the distribution. Red lines: percent of la-EPC loci in the same cell line with an overlapping CTCF ChIP-seq peak. **B**. The la-EPC preference scores of enrichment of ChIP-seq peaks at la-EPCs compared to na-EPCs. Significantly enriched and depleted proteins (for which data exists in at least two cell lines, with an empirical p value < 0.05 in all cell lines for which data is available) are highlighted in red and green, respectively. **C**. Correlation between K562 ChIP-seq–based la-EPC preference scores and the percent of ChIP-seq peaks which overlap protein-coding gene TSS. Spearman correlation coefficient and p value are shown. Coloring indicates local point density. **D**. Distribution of average la-EPCs preference scores in K562, HepG2, and MCF7, grouped by Interpro protein domain annotations. P values calculated using two-sided Wilcoxon test; Interpro domains associated with p values < 0.05 are indicated by red asterisks. E. ChIP-seq–based la-EPC preference scores for proteins with DNA binding motifs enriched or depleted in la-EPCs vs. na-EPCs. P values calculated using two-sided Wilcoxon test. **F**. G/C content of regions ±1Kb around PCG TSSs, la-EPCs, and na-EPCs from the indicated cell lines. P values calculated using two-sided Wilcoxon test.

For each protein profiled in the ENCODE datasets, we calculated the “la-EPC preference” score, which is the Z score of the fraction of ChIP-seq peaks overlapping la-EPCs compared to random sets of H3K27ac-matched na-EPCs (see Methods). la-EPC preference scores are highly correlated between the different cell lines (Figure S3B), suggesting that la-EPCs are associated with consistent differential protein binding across cell lines. 154 proteins, including TFs such as JunD, SP5, Klf9, GABPA, and Myc, as well as chromatin modifiers associated with RNA-dependent modulation such as WDR5 and BMI1 (Yang et al. 2014; Hu et al. 2014), are significantly associated with la-EPCs in all cell lines for which data is available (Figure 3B; **Table S2**). Interestingly, Ago1, which has been reported to bind eRNAs (Alló et al. 2014), was also found to be significantly enriched in la-EPCs in the one available dataset.

We next hypothesized that proteins that preferentially bind la-EPCs compared with na-EPCs would also be preferentially found at PCG promoters, as la-EPC regions serve as promoters for lncRNA genes. Indeed, there is a strong correlation between the la-EPC preference score and the fraction of ChIP-seq peaks of that protein which are associated with PCG promoters (**Table S2**; Figures 3C and S3C). lncRNA-producing enhancers therefore harbor promoter characteristics, including a distinct protein-binding landscape. Examination of domains enriched in proteins with high la-EPC preference scores identified specific families, including basic helix-loop-helix domains, which are present in many dimeric TFs (Jones 2004) such as Myc (**Table S2**; Figure 3D). Interestingly, there is also a significant enrichment for RNA Recognition Motif (RRM) domains at la-EPCs, while proteins preferentially bound at na-EPC are enriched with OB-fold nucleic acid-binding domains (Figure 3D), suggesting distinct nucleic acid-binding domains are associated with establishment of enhancer regions with different propensity for producing stable RNAs.

Interestingly, proteins containing Forkhead domains are significantly depleted at la-EPCs compared to na-EPCs (Figure 3D). FOX proteins act as pioneering TFs that are able to bind to condensed chromatin, which in turn enables the recruitment of additional TFs and the Pol II preinitiation complex (Lalmansingh et al. 2012; Cirillo et al. 2002) (see Discussion).

### lncRNA-associated EPCs are associated with specific TF binding motifs

Despite the positive correlation between lncRNA transcription and enhancer activity, the causality remains undetermined: are more active enhancers able to generate lncRNA transcripts due to their higher Pol II recruitment and transcription levels, or does lncRNA production cause an enhancer to be more active, perhaps through the recruitment of specialized factors to the enhancer locus? In order to differentiate between these two possibilities, we used Analysis of Motif Enrichment (AME) (McLeay and Bailey 2010) to compare the sequences of the two sets of EPCs and their flanking regions. In all three cell lines tested, there is a correlation between motif enrichment and the ChIP-seq-derived la-EPC preference scores for the corresponding factors (**Table S2**; Figure 3E), suggesting that the differential protein occupancy at la-EPCs is encoded in the DNA sequence of these loci. This also affirms that the protein binding at la-EPCs is not the result of indirect or “phantom” peaks due to long-range interactions between the enhancers and PCG promoters (Liang et al. 2014; Jain et al. 2015). Our identification of specific proteins/binding motifs which are preferentially associated with enhancer-only or with both promoter and enhancer activities is consistent with focused studies of specific sequences in mouse neurons (Nguyen et al. 2016), which identified some of the same motifs – including binding sites for GABPA, EGR1, and KLF – as being enriched in regions displaying high promoter activity. Similarly, the flanking G/C content of la-EPCs is intermediate between that of PCG TSSs and na-EPCs (Figure 3F), in agreement with previous findings (Robin Andersson, Gebhard, et al. 2014).

### Increased splicing at the lncRNA region is associated with higher accessibility at regions flanking the EPC

Among the proteins with strong la-EPC preference scores in the ChIP-seq data are several proteins involved in RNA splicing, including RBFOX2, SRSF1, U2AF1, and TARDBP (**Table S1**). lncRNAs, unlike eRNAs, are usually spliced (De Santa et al. 2010; Ulitsky and Bartel 2013), as are >90% of the lncRNAs that are associated to an EPC in our datasets; we therefore examined the association between lncRNA splicing and enhancer activity. In lncRNAs associated to EPCs we noted a strong correlation (R=0.66; Figure 4A) between the density of the exons and the density of the DHSs (density, which is calculated by dividing the number of exons/DHSs by the locus length, cancels out the effect of the length; Figure S4A). Further, exon density is also weakly correlated with the H3K27ac levels of the DHSs found within the lncRNA sequence (Figure 4B). Together, these results suggest that more densely spliced lncRNAs are associated with more active enhancers. Interestingly, we did not see a similar correlation between the lncRNA exon density and the H3K27ac levels in a small window around the EPC itself (±1Kb, R=-0.0215), suggesting that splicing is associated with the activity of the broader enhancer region rather than the focal region of the lncRNA promoter.

**Figure 4.**
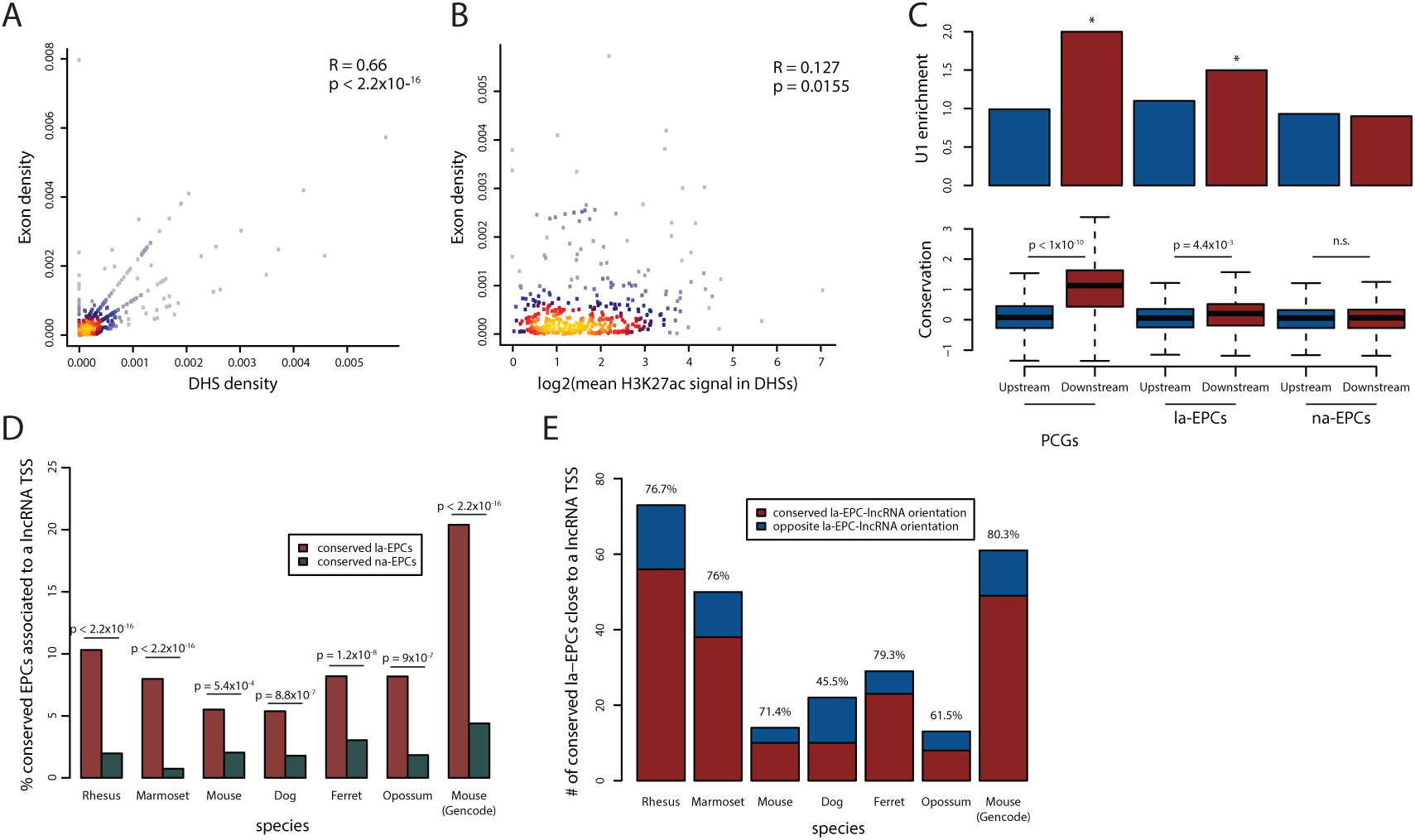
Conservation of la-EPCs. **A**. Density of exons and density of DHSs within the regions of K562 lncRNAs associated to EPCs. Spearman correlation coefficient and p value are shown. Coloring indicates local point density. **B**. Density of exons and mean H3K27ac signal in the regions ±200bp around DHSs found within K562 lncRNAs that are associated to EPCs. Spearman correlation coefficient and p value are shown. Coloring indicates local point density. **C**. Occurrence of the U1 splicing motif (top) and PhyloP conservation signal of the motifs (bottom) in the indicated regions ±500bp from PCG TSS, la-EPCs, or na-EPCs. Regions from K562, HepG2, and MCF7 cells were combined. **D**. Percent of the regions aligning to human EPCs of the indicated type that are located <500bp from a lncRNA TSS in the indicated species. Proportion test p values are shown. **E**. Conserved la-EPCs that are also associated to lncRNAs in the indicated species, split based on the orientation of the lncRNA transcript relative to the EPC sequence (which is based on the human EPCs). Numbers above bars are percentage of lncRNA-EPC pairs in the indicated species with the same relative orientation as in human.

The increase in DHS density associated with higher exon density could be a direct result of lncRNA transcription/processing that may affect chromatin accessibility, or it could indirectly reflect remodeling of the broader region. To distinguish between these possibilities, we compared the chromatin landscape at EPC-associated lncRNA gene body regions with size-matched regions on the other side of the EPC. Intriguingly, there is a strong correlation between the number of DHSs on both sides of the EPC (on the one where the lncRNA is produced, and on the matching side which does not produce a lncRNA) (Figure S4B). The DHSs on the two sides of the EPC are also similar in terms of their average H3K27ac signal (Figure S4C). This indicates that the broader enhancer region is more active when a lncRNA is transcribed and spliced at that enhancer. This “ripple” effect points at recruitment of activating factors to the locus by either the lncRNA or the act of its transcription/maturation as the driver of higher enhancer activity, rather than a local effect of Pol II elongation.

### Sequence-encoded U1 binding sites with signs of evolutionary conservation dictate the direction of lncRNA production from the EPCs

The production of spliced lncRNAs at la-EPCs and not at na-EPCs could either be encoded in the DNA sequence of the loci or be the result of the differential protein binding landscape. To distinguish between these options, we next examined the DNA sequence at the two sides of the la-EPCs, and compared them to PCG TSSs and na-EPCs. As noted previously (Almada et al. 2013), the U1 splicing motif is enriched downstream of PCG TSSs to the direction of the mRNA transcription (Figure 4C, **top**). A similar enrichment was observed at la-EPCs, only in the direction of lncRNA production, but not around na-EPCs. Importantly, the U1 sites found downstream to PCG TSSs and la-EPCs also exhibit significant signals of sequence conservation compared to those found on the other side of the TSS (Figure 4C, **bottom**), suggesting that natural selection acts on the splicing signals that allow the formation of spliced lncRNA transcripts at some enhancers.

### Production of lncRNAs at la-EPCs is conserved in evolution

In order to analyze conservation of lncRNA production at la-EPCs, we projected regions of human EPCs to genomes of six other mammals (see Methods). Regions aligning to human la-EPCs in other species tend to have a lncRNA TSS within several hundred bases (Figure S4D), and a significantly higher percentage of regions aligning to an la-EPC compared to those aligning to an na-EPC are located nearby (<500bp) a lncRNA TSS (Figure 4D) in all species examined, indicating that the ability of specific enhancers to drive lncRNA formation is a conserved feature of these enhancers. The exact number of EPCs that are associated with lncRNA transcription in another species is naturally heavily dependent on the lncRNA annotations, as evident when using the comprehensive GENCODE annotations vs. PLAR annotations that are based on one RNA-seq dataset (Hezroni et al. 2015) (Figure 4D). Interestingly, in some cases where there is a lncRNA in another species, its orientation relative to the enhancer is different from their relative orientation in human (Figure 4E). This might indicate that the actual act of lncRNA transcription is more important for enhancer activity than the lncRNA sequence or even the direction of its production.

## Discussion

The diversity of transcripts produced at enhancer elements obscures the biochemical and functional distinctions between enhancers and promoters. Here, we show that some genomic regions with enhancer characteristics serve as promoters for the production of lncRNA transcripts. These regions are associated with stronger enhancer activity, which is encoded in their sequences (Figure 2D) and appears to be driven in part by the presence of conserved, directional splicing signals that drive lncRNA production. Appearance in evolution of lncRNA transcripts in intergenic regions is associated with modestly higher expression of the flanking PCGs (Kutter et al. 2012); consistently, recent studies, based on methods complementary to the ones we used here, reported that splicing of RNA transcripts, both coding and noncoding, could increase the expression of nearby genes (Engreitz et al. 2016; Tan et al. 2018). Together with our findings, the emerging conclusion is that RNA splicing and/or recruitment of the spliceosome plays a general role in sculpting the local chromatin/activity state of the underlying genomic region, thereby causing enhanced expression of the associated PCGs.

It is presently unclear through which mechanism splicing could affect the activity level of the underlying enhancer. However, this effect is not likely to be restricted to the boundaries of the lncRNA gene body or its splice sites, as stronger enhancer activity appears distributed throughout the region and is not limited to the side of the EPC to which the lncRNA is transcribed (Figures S4B,C). One possibility is that splicing factors, once recruited to the locus, engage with additional proteins that remodel the enhancer, for example through chromatin modifications (Schüler, Ghanbarian, and Hurst 2014). Another possibility is that splicing functions indirectly, for example through increasing the efficiency of transcription elongation (Fong and Zhou 2001; Brinster et al. 1988), likely through interactions between splicing proteins and the transcriptional machinery (Hirose, Tacke, and Manley 1999; McCracken et al. 1997), and that the resulting accumulation of Pol II and its associated proteins leads to the recruitment of factors that increase enhancer activity.

Another particularly appealing possibility is that lncRNA transcription and splicing affect enhancer activity through generally “opening” the locus or maintaining it at an open conformation. For example, splicing could promote the dissociation of the lncRNA from the chromatin, enabling the binding of additional factors, as has been proposed for the A-ROD lncRNA (Ntini et al. 2018). This general open state of the enhancer could then affect its position within the nucleus (Isoda et al. 2017) or the rate of locus mobility in the nucleus (Gu et al. 2018). This model is supported by the strong correlation between the exon and DHS density in la-EPCs (Figure 4A). Intriguingly, la-EPCs and na-EPCs are bound by proteins with different characteristic nucleic acid-binding domains; the preferential recruitment of proteins containing FOX domains, found in several pioneering TFs, to na-EPC regions (Figure 3D) suggests that la-EPC regions may employ alternative methods for opening the chromatin and initiating transcription, possibly pointing at a pioneering role for the lncRNA production.

Whatever the mechanism, so far there is limited evidence that the RNA products of lncRNAs produced at EPCs play a role in enhancer activity. The sequences of enhancer-associated lncRNAs show virtually no signs of conservation (Marques et al. 2013), and when comparing mammals and other vertebrates, most lncRNAs appear to be found in syntenic regions without any detectable sequence alignability (Hezroni et al. 2015). The scarce conserved sequence motifs in lncRNAs tend to be associated with splicing (Schüler, Ghanbarian, and Hurst 2014; Haerty and Ponting 2015). Relatedly, as we show here, the direction to which the lncRNA is transcribed from the EPC is not always conserved between mammals (Figure 4E), and is not associated with any of the examined features of enhancer activity. There is also no preferential directionality for CTCF DNA motifs at la-EPCs, which are overall enriched for CTCF binding (Figure 3A), relative to the direction of the lncRNA transcription (data not shown). The specific exon-intron architecture of these lncRNAs is also typically poorly conserved (this ‘structural’ evolution is more difficult to quantify, since individual lncRNA exons are typically not readily alignable between human and non-primate species) (Ulitsky 2016). All these suggest that the act of transcription, coupled to recruitment of the splicing machinery, constitute the functionally relevant aspects of lncRNA production from enhancer regions.

It is interesting to consider our results in the context of evolution of new lncRNA genes. New lncRNAs can form through a variety of scenarios (Ulitsky 2016), including duplication of existing lncRNAs, pseudogenization of PCGs, and exaptation from DNA regions that did not produce stable transcripts. We have previously shown that evolution of new lncRNAs by duplication or pseudogenization likely occurs quite rarely (Hezroni et al. 2015, 2017). Thus, the more common scenario for the evolution of a new lncRNA gene involves the coming together of a promoter, splice sites, and/or poly(A) sites, contributed mostly by random mutations or from transposable elements. The fact that enhancers recruit Pol II and associated factors, and that this recruitment results in eRNA transcription, makes them an attractive source of promoters for new lncRNAs.

Indeed, it has been observed that promoters of some mouse lncRNAs expressed in mES cells map to human regions with chromatin marks indicative of enhancer activity but no lncRNA production (Engreitz et al. 2016). Furthermore, Wu and Sharp proposed a model in which increased transcription in the germline leads to increase in mutations to G/T, which strengthens the U1-PAS axis, constituting a positive feedback loop which drives the formation of new genes divergent to existing PCG promoters (Wu and Sharp 2013). Similar evolutionary trajectories are likely to happen at enhancer regions, where mutations that lead to the gain of binding of TFs that favor lncRNA production can lead to enhanced transcription, and strengthening of the U1 axis, favoring production of processed and stable transcripts.

Our results suggest two possible models: i) some enhancers, whose protein binding landscape resembles that of PCG promoters, are more common substrates for evolution of new lncRNAs; or ii) once a combination of events leads to lncRNA production from an enhancer, subsequent evolution introduces binding sites for proteins that are typically associated with promoters and which may increase enhancer activity. Distinguishing between these two models will be challenging, as it will require data that allows comparing la-EPCs and na-EPCs in similar cell types across several species, alongside information on the proteins that are binding each region. Some relevant data are now beginning to become available (Danko et al. 2018), but large-scale, directly comparable, ChIP-seq data are presently available only for few human cell types.

The ability to produce a mature lncRNA transcript thus distinguishes a group of enhancers with higher activity, which appears to be dependent at least in part on the processing events involved in lncRNA maturation. Detailed and large-scale perturbations of endogenous loci producing individual lncRNAs, which are becoming more accessible using CRISPR/Cas9 technologies, should allow us to increase our understanding of how signals encoding lncRNA production affect enhancer functionality.

## Methods

### Datasets

| Description | Downloaded from / accession | Cell types |
| --- | --- | --- |
| DNAse hypersensitive sites | ENCODE | K562, HepG2, MCF7, mES |
| GRO-seq | GSE60454, GSE92375, GSE43835, GSE69142 | K562, HepG2, MCF7, mES |
| H3K27ac ChIP-seq | GSM733656 , GSM733743, GSM945854, GSM1000126 | K562, HepG2, MCF7, mES |
| H3K4me1 ChIP-seq | GSM733692, GSM798321, GSM2308526, GSM1003750 | K562, HepG2, MCF7, mES |
| Chromatin segmentations | ENCODE | K562, HepG2 |
| RNA-seq | GSM958731, GSE90322, GSE78663, GSE93453 | K562, HepG2, MCF7, mES |
| Pol II ChIA-PET | GSE33664 | K562, MCF7 |
| Pol II ChIA-PET | GSM1084138 | mES |
| YY1, CTCF ChIA-PET | GSE99520 | mES |
| STARR-seq | GSE100432 | HeLa |

### EPC identification

Regions of bidirectional transcription were identified by searching for bidirectional GRO-seq read coverage in a region ±2Kb away from DHSs, as follows (Figure S1A): We iterated over the DHS regions and computed, for each DHS, the GRO-seq reads in the region between the DHS start and 2Kb downstream of its end (‘plus_signal’), and from 2Kb upstream of its start to the DHS end (‘minus_signal’). Only DHSs with min(plus_signal,minus_signal)>=2 (>=8 for the mES dataset) were considered further. In those regions we found the EPCs by computing a ‘separation’ score for each position in the DHS region padded by 2Kb on each side. The separation score for position *P* was defined as the geometric mean of (a) the fraction of the GRO-seq signal on the minus strand to the left of *P* out of *minus_total*, and (b) the fraction of the GRO-seq signal on the plus strand to the right of *P* out of *plus_total*. The position achieving the highest score (P_max_) was defined as the EPC. If multiple genomic positions had the same P, the midpoint of those positions was defined as the EPC. To compute eRNA boundaries, we identified P_first_ as the leftmost position within the considered region where 90% of the *minus_total* signal is found to the right of it, and P_last_ as the rightmost position within the considered region where at least 90% of the *plus_total* signal is found to the left of it. The regions (P_first_,P_max_) and (P_max_,P_last_) were considered as the eRNA pair. In case of overlapping pairs, only the pair with the higher minimal *sum* of GRO-seq signal on the two sides of the EPC was retained.

Only EPCs located >10Kb away from the nearest RefSeq PCG were considered in the following analyses. la-EPCs and na-EPCs were classified as EPCs located <200/300bp (human/mouse) and >3Kb, respectively, from the nearest TSS of a GENCODE lncRNA. A combined list of EPCs from all three human cell lines was prepared by selecting the EPC with the highest signal sum in each window of 200bp.

### Analysis of EPCs and flanking regions

For intersection of EPCs with ENCODE chromatin segmentations, the ENCODE combined segmentation data (Hoffman et al. 2013) for the cell lines K562 and HepG2 was downloaded. Overlap of the EPC with each of the chromatin states was calculated using BEDTools (Quinlan and Hall 2010). Percent overlap with enhancer states (‘Enhancer’ or ‘Weak enhancer’) was computed using the BEDTools window tool, with a 1Kb window around the EPC.

The groHMM algorithm (Chae, Danko, and Kraus 2015) was used to annotate MCF7 eRNAs from the same GRO-seq dataset used in this study, employing parameters used in the original manuscript (LtProbB=-200, UTS=5, threshold=1). Resulting transcripts were selected for bidirectional transcripts (overlapping at any point on opposite strands) whose TSSs were >10Kb from the nearest PCG. These eRNA were compared to our set of MCF7 EPCs by quantifying the overlap between regions ±2Kb from MCF7 DHSs and the eRNAs.

Metagene plots were prepared using the ngs.plot module (Shen et al. 2014) with the following parameters: −L 4000, −RB 0.02. Signals from BAM files (see table of datasets used in this study) were centered around the EPCs.

Quantification of chromatin marks was performed by taking the mean H3K27ac signal ±1Kb around EPCs, or ±200bp around DHSs, extracted from the bigwig files using the bigwig_file module from the python bx.bbi package.

Overlap between EPCs and anchors of significant Pol II ChIA-PET interactions was calculated using BEDTools. For mES ChIA-PET analysis, la-EPC and na-EPC coordinates were first converted to mm9 using UCSC liftOver tool (Hinrichs et al. 2006).

### Protein binding enrichments

All available BED files containing ChIP-seq peaks for each of the cell lines were downloaded from the ENCODE website (K562/HepG2 data: October 2017, MCF7/mES data: March 2018). BEDTools was used to compute overlap between each of the BED files and either la-EPCs or na-EPCs. la-EPCs were divided into five bins according to their H3K27ac levels ±1Kb around the EPC. Percentages of overlap of the ChIP-seq peaks with na-EPCs were calculated in 1000 random samples of na-EPCs with H3K27ac levels matched to those of the la-EPCs. The distribution of the percent overlap of each protein over the na-EPCs was plotted, and the empirical p value was computed. The “la-EPC preference” score, which is the Z score of enrichment/depletion of ChIP-seq peaks in the la-EPCs vs. the random set of the na-EPCs, was calculated by subtracting the mean percent of na-EPCs with an overlapping ChIP-seq peak (over the 1,000 random samples) from the percent of la-EPCs with an overlapping ChIP-seq peak, and dividing by the standard deviation of the na-EPC sampling distribution.

Overlap of ChIP-seq peaks of each protein with RefSeq PCG TSSs was calculated using BEDTools. A list of the Interpro domains of each protein was downloaded from the Ensembl site.

### Motif enrichment and sequence composition in la-EPCs vs. na-EPCs

Analysis of Motif Enrichment (AME) (McLeay and Bailey 2010) was used to determine motif enrichments/depletions in sequences ±2Kb around la-EPCs, na-EPCs, and RefSeq PCG TSSs.

G/C content for the sequences in the regions ±2Kb around la-EPCs, na-EPCs, and PCG TSSs was calculated using the Biostrings package (Pages et al. 2016).

### U1 motif enrichment and conservation

For U1 motif analysis, we extracted the genomic sequences in regions of 0.5Kb flanking the EPCs, and counted the number of occurrences of one of three common U1 sequence motif variants (Almada et al. 2013) – GGTAAG, GGTGAG, and GTGAGT. For na-EPCs, direction of transcription of the nearest lncRNA was chosen as reference. For enrichment analysis we compared the number observed in 1,000 randomly shuffled sequences of the same length. The shuffling was performed by dividing the sequence into windows of 100bp each, and separately shuffling each window while presenting dinucleotide composition. For each match to the motif, we computed the average PhyloP score (Pollard et al. 2010) in a 100-way whole genome alignment (scores obtained from the UCSC genome browser). In order to normalize the enrichment of the motif to the background conservation in the entire region, we then Z-transformed the enrichment using the mean and the standard deviation of the PhyloP scores in the examined region (EPC±0.5Kb).

### EPC conservation

Regions ±30bp around human la-EPCs and na-EPCs were converted to rheMac3 (rhesus macaque), calJac3 (marmoset), canFam3 (dog), mm9/mm10 (mouse), musFur1 (ferret), and monDom5 (opossum) genome assemblies using UCSC liftOver tool (Hinrichs et al. 2006). lncRNA TSS coordinates were extracted from tables derived by PLAR (Hezroni et al. 2015) for all species except mm10, for which GENCODE vM16 lncRNA TSSs were used. Closest lncRNA TSS was identified using BEDTools; EPCs were considered to be lncRNA-associated in the other species if they were found <500bp from a lncRNA TSS. The conservation of the direction of lncRNA transcription was examined by comparing the strand to which the EPC was mapped in the other genome with the direction of the transcription of the associated lncRNA.

## Supplementary Figures

**Figure S1.**
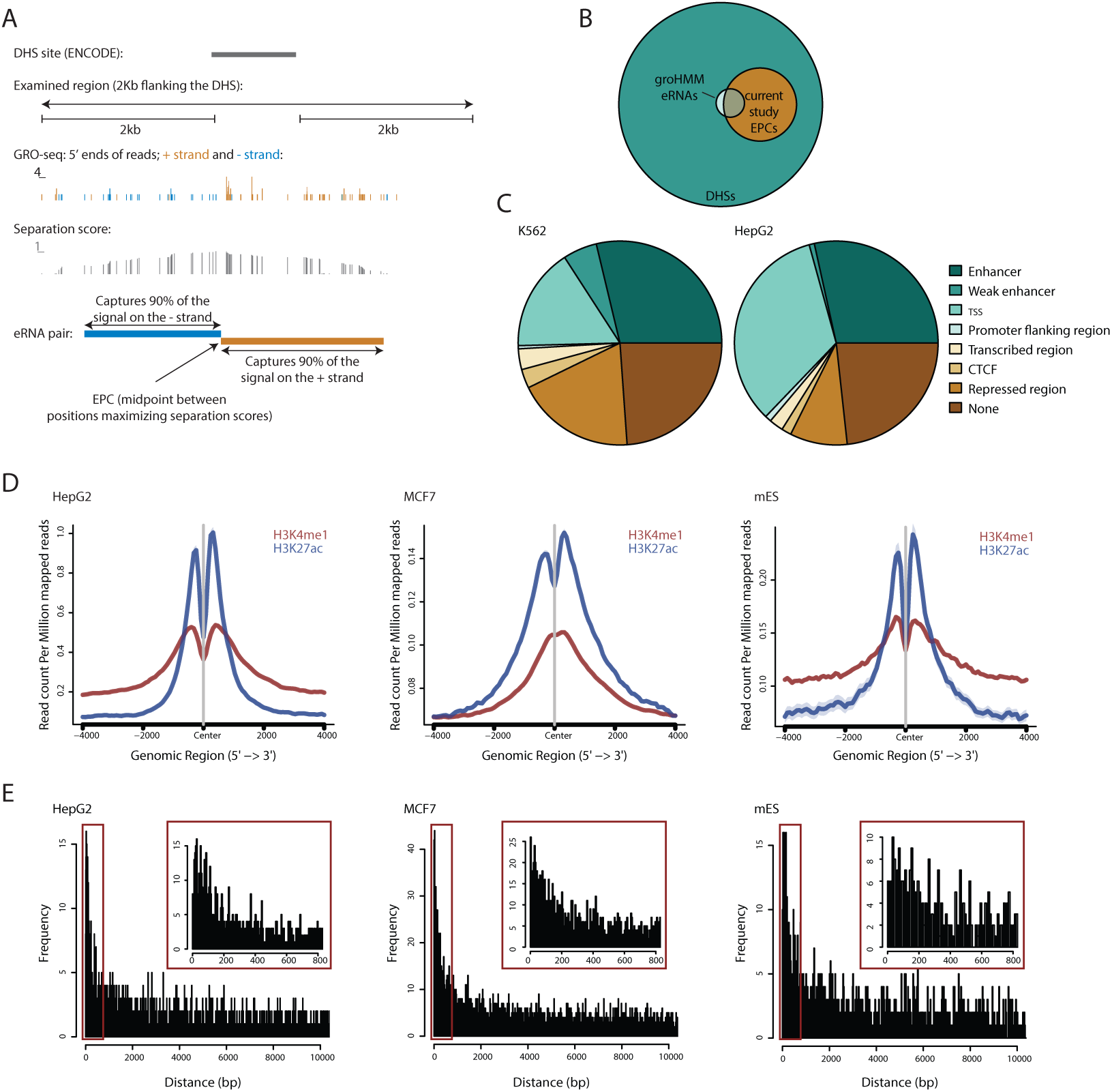
**A**. Genome browser image for the hg19 coordinates chr17:79,452,917-79,457,070, with an example of the scheme used for annotating EPCs from DHS data (see Methods). **B**. An Euler diagram showing overlap between DHSs and the indicated groups of transcripts in MCF7 cells (see Methods for details of groHMM execution). **C**. Intersection of EPCs and chromatin segmentation annotations (obtained from ENCODE) in the indicated cell lines. **D**. Metagene plots showing distribution of H3K27ac and H3K4me1 ChIP-seq signal around the EPCs from the indicated cell lines. Shaded regions represent standard errors. **E**. Distribution of distances between EPCs and the closest GENCODE lncRNA TSS for the indicated cell lines. Regions in the red rectangles are shown with higher bin resolution in insets.

**Figure S2.**
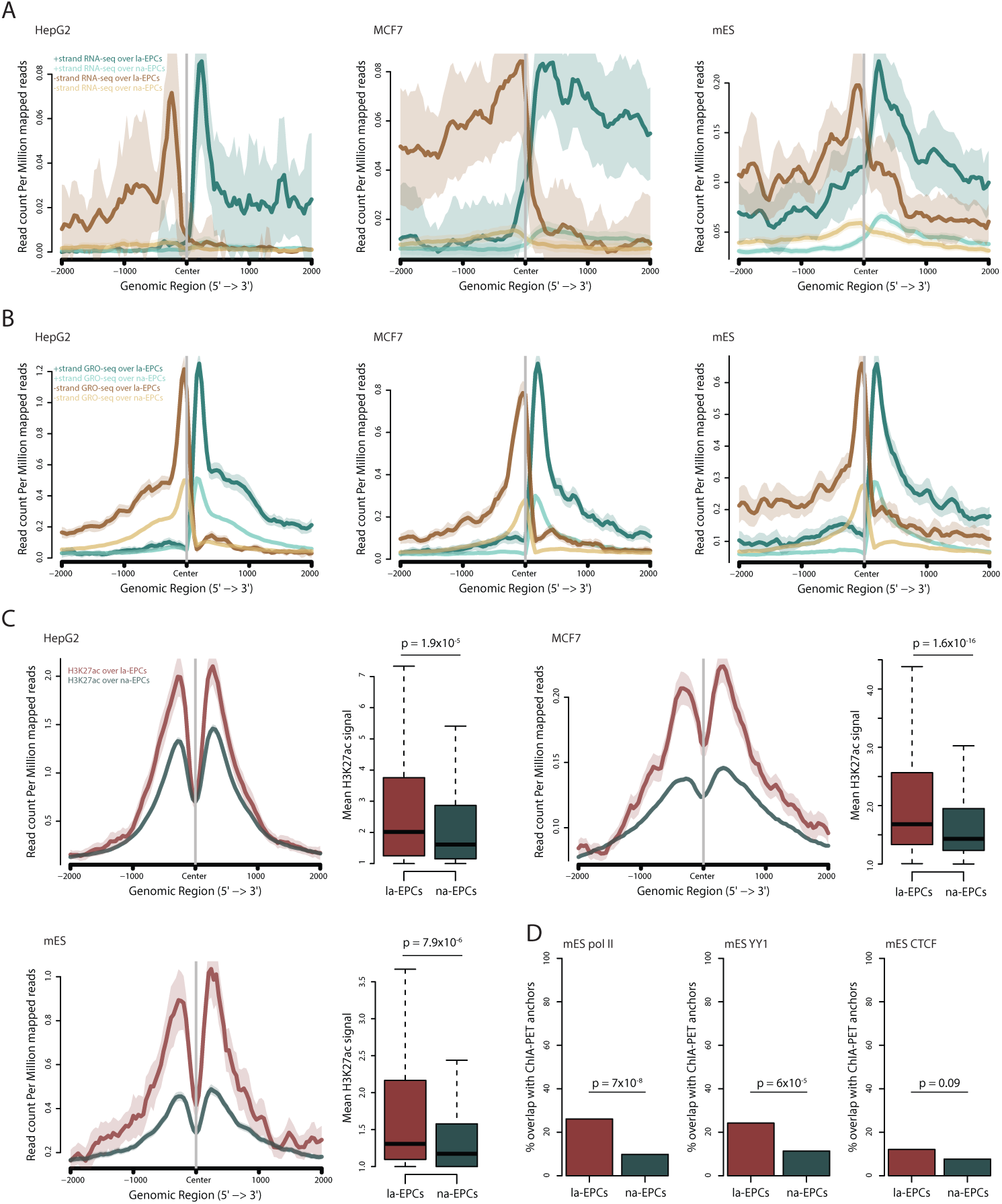
**A**. Metagene plots of RNA-seq signal separated by strand from the indicated cell lines over la-EPCs and na-EPCs. Shaded regions represent standard errors. **B.** As in (A), for GRO-seq signal. **C.** Metagene plots of H3K27ac ChIP-seq signal from the indicated cell lines over la-EPCs and na-EPCs, alongside quantification of H3K27ac signal over the region ±1Kb around EPCs from the indicated cell lines. P values were calculated using two-sided Wilcoxon test. Shaded regions represent standard errors. **D.** Percent of mES la-EPCs or na-EPCs which overlap Pol II/YY1/CTCF ChIA-PET anchors. P values calculated using the proportion test.

**Figure S3.**
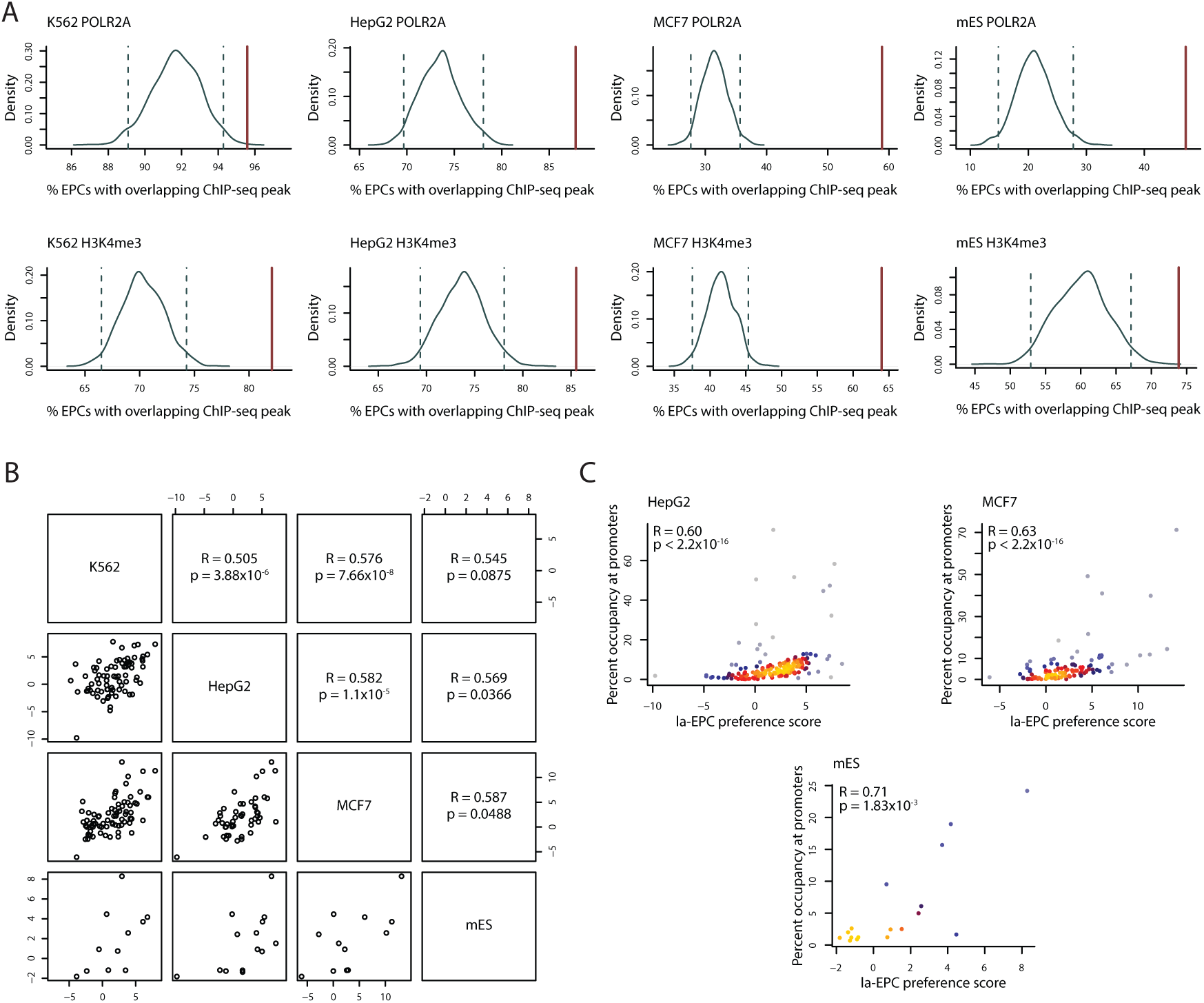
**A**. Green lines: distribution of the percent of na-EPC loci with overlapping Pol II (top) or H3K4me3 (bottom) ChIP-seq peaks in a random sampling of na-EPC loci in the indicated cell lines. Dashed lines represent 2.5^th^ percentiles of the distributions. Red lines: the percent of la-EPC loci in the same cell line with overlapping ChIP-seq peaks. **B**. Correlation between la-EPC preference scores in the different cell lines used in this study; each dot represents the la-EPC preference score of a single protein. **C**. Correlation between la-EPC preference in the indicated cell line, and the percent of ChIP-seq peaks which overlap protein-coding gene TSSs. Spearman correlation coefficients and p values are shown. Coloring indicates local point density.

**Figure S4.**
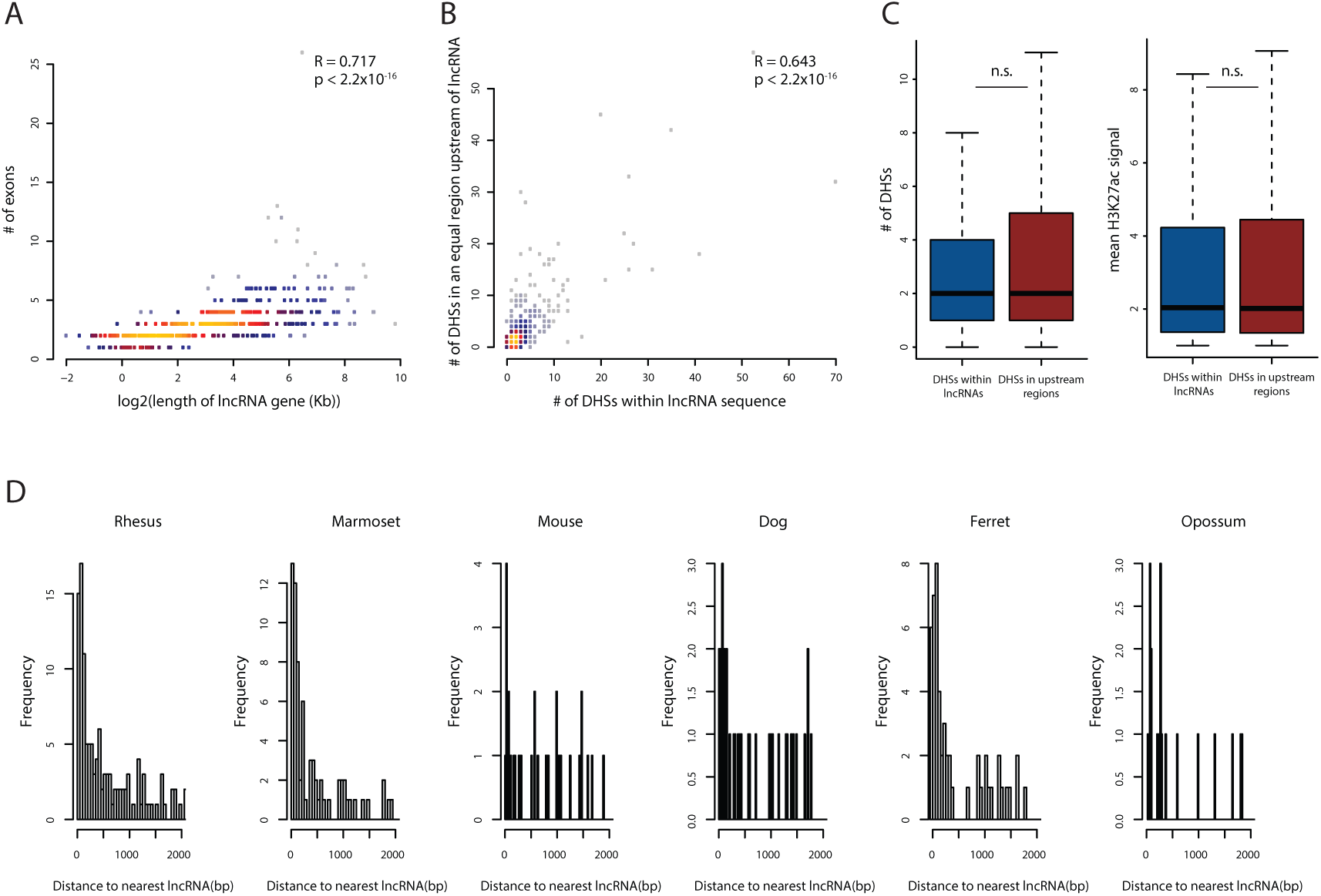
**A**. Total length of lncRNAs associated to EPCs and the numbers of exons within their sequence. Spearman correlation coefficient and p value are shown. Coloring indicates local point density. **B**. Correlation between the number of DHSs on the two sides of la-EPCs, within regions equal to the locus length of the associated lncRNA. Spearman correlation coefficient and p value are shown. Coloring indicates local point density. **C**. Number of DHSs (left) or mean H3K27ac ChIP-seq coverage at the DHSs (right) found on the two sides of la-EPCs, within regions equal to the locus length of the associated lncRNA. P values calculated using two-sided Wilcoxon test. **D**. Distribution of distances between la-EPCs projected from human and the closest lncRNA TSS in the indicated species.

**Supplemental Table 1 key**. Each sheet contains hg19/mm10 EPC coordinates from the indicated cell line, as well as their distance to the nearest GENCODE lncRNA TSS and classification into la-EPCs or na-EPCs (related to Figures 1A and S1A).

**Supplemental Table 2 key**. For each protein the la-EPC preference score in the indicated cell lines is provided, as well as indication of significance (related to Figures 3A,B and S3A,B); the percent of ChIP-seq peaks of the protein which overlap PCG TSSs in the indicated cell lines (related to Figures 3C and S3C); the Interpro Short Description of the domains associated with the human protein (related to Figure 3D); and an indication of whether the DNA motifs associated with the protein are significantly enriched/depleted in the indicated cell lines (related to Figure 3E).

